# Transcriptomic Insights into Drought Tolerance Enhancement in Bread Wheat Induced by a Microalgae-based Biostimulant

**DOI:** 10.64898/2026.05.18.725825

**Authors:** Christina Arvanitidou, Marcos Ramos-González, M. Elena García-Gómez, Mercedes García-González, Francisco J. Romero-Campero

## Abstract

Bread wheat (*Triticum aestivum*) is a staple food crucial for global caloric intake and food security. The current climate emergency demands the development of sustainable agricultural practices, particularly in the context of drought-induced yield reductions in bread wheat. Microalgae-based biostimulants have emerged as promising tools to enhance crop tolerance to drought stress while concurrently mitigating atmospheric CO_2_ accumulation. This study characterizes the transcriptomic responses to the foliar application of the microalgae-based biostimulant LRM^TM^ in drought-stressed and fully irrigated wheat plants unveiling its mode of action. Drought stress at the tillering stage significantly altered gene expression activating key pathways related to phosphate starvation response (PSR), inositol phosphate signaling, and tocopherol biosynthesis. The application of the microalgae-based biostimulant LRM^TM^ in drought-stressed plants further enhanced the expression of drought-responsive genes, particularly those involved in PSR and carbon fixation. Specific responses to LRM^TM^ treatment in drought-stressed plants were also found related to abscisic acid (ABA) signaling activating genes involved in stomata closure, which plays a critical role in drought tolerance. In fully irrigated plants, LRM^TM^ treatment was also beneficial modulating circadian rhythms, shade avoidance and attenuating stress responses. Phenotypic analysis showed that LRM^TM^-treated plants exhibited enhanced drought tolerance, increased height and spike length even under fully irrigated conditions. These results indicate that the microalgae-based biostimulant LRM^TM^ not only enhances wheat response to drought but also promotes growth and productivity in both stressed and non-stressed conditions which could contribute to the development of sustainable agriculture in the face of the current climate challenges.

## Introduction

The steady increase in world population, coupled with the current climate emergency, makes the development of new sustainable agricultural practices imperative. These new methods are necessary to enhance crop tolerance to the adverse conditions produced by global warming, such as drought, while increasing crop yield at the same time (Ronga et al., 2019). Agricultural consumption is expected to increase by 60% in 2050. This rise in demand cannot be sustainably addressed by expanding the use of chemical fertilizers due to their harmful impact on soil composition and the carbon footprint associated to their production. Specifically, chemical fertilizers have been shown to alter the nitrification process, with effects on pH and eutrophication (Ayiti & Babalola, 2022; Chabili et al., 2024; FAO, 2018). Moreover, climate change caused by atmospheric CO_2_ accumulation as a result of human activity is estimated to produce crop losses in the range of 30-70%. Therefore, it becomes crucial to address these challenges by developing novel sustainable agricultural stimulants whose production process is coupled with atmospheric CO_2_ abatement.

In this respect, biostimulants derived from microalgae extracts are emerging as promising plant growth promoters and crop stress alleviators since microalgae biomass production is coupled with CO_2_ fixation (Arvanitidou et al., 2024; Chabili et al., 2024; Kapoore et al., 2021). In addition, microalgae biomass can be produced in a cost-effective way by applying the principles of circular economy, using waste products from human activities, such as wastewater or anaerobic digestion products (Chiaiese et al., 2018; Fernández-Rodríguez et al., 2022; Kapoore et al., 2021). However, traditionally, macroalgae (seaweeds) biomass has been used as the raw material for the development of agronomic products, with interest in microalgae arising only recently (Arvanitidou et al., 2024; Chabili et al., 2024). Among the most widely used seaweeds, stramenopile macroalgae such as *Ascophyllum nodosum* and *Durvillaea potatorum* have been shown to confer agricultural desirable traits such as flowering anticipation or resistance to abiotic stresses (H. S. S. Sharma et al., 2014). A potential issue with using seaweed for biostimulants production is that they are predominantly wild harvested. This raises concerns about the preservation of ecosystems and their biodiversity, which has led to regulations restricting the use of specific macroalgae species and their collection in certain areas (Kapoore et al., 2021). More importantly, wild harvesting implies that macroalgae are subjected to climatic and seasonal changes that affect the composition of their extracts, preventing consistency in their effects.

In contrast, microalgae are successfully cultivated at industrial levels in confined and fully controlled conditions, allowing for the monitoring and fine-tuning of growth parameters such as growth media, temperature and illumination regimes. These methods ensure a desirable composition of the microalgae extracts, solving the previously mentioned consistency issue in macroalgae extracts (Chiaiese et al., 2018). Another important difference regarding their composition is the higher protein content in microalgae extracts when compared to their macroalgae counterparts, reaching 18-46% of dry weight (Chiaiese et al., 2018). These molecules are particularly relevant in crop biostimulation since amino acids are precursors of many plant phytohormones (Malécange et al., 2023).

Regarding the evolutionary history of micro/macroalgae and crops, land plants, or embryophytes, including crops, constitute together with chlorophytes and streptophytes, the green lineage or Viridiplantae. This group diverged from stramenopiles around two billion years ago. Therefore, chlorophytes and streptophytes green microalgae share many biochemical and signaling systems with crops, making extracts from these organisms elicit responses in land plants. In contrast, the systems in stramenopile macroalgae are markedly different from those in crops, which is expected to render plants less sensitive to macroalgae extracts (Bachy et al., 2022; Strassert et al., 2021). These advantages are shifting interest from macro to microalgae derived products for agronomic applications (Chiaiese et al., 2018).

Bread wheat (*Triticum aestivum*) represents the second cereal crop in terms of global production, providing a significant portion of the caloric intake for human population and ensuring food security (FAO, 2018). Bread wheat is rich in carbohydrates, provides essential proteins and contributes vitamins and minerals to the human diet. Its predominant role as staple food is mainly due to its adaptability to a wide range of temperate climates together with its high nutritional value (Shewry, 2009). Nonetheless, the current climate emergency, characterized by rising global temperatures and increasing frequency of droughts, is leading to significant reductions in grain yield and quality in bread wheat. Reduced water availability affects wheat plant growth at all developmental stages, from tillering to flowering and grain-filling, which are critical for determining the final yield (I. Sharma et al., 2015). The development of innovative agricultural practices, such as the one proposed here based on microalgae extracts, is essential to address the challenges imposed by these changing climatic conditions to sustain and enhance wheat production (Farooq et al., 2024).

Despite efforts to optimize production and characterize the composition and effect of microalgae-based biostimulants on crops, few studies have focused on detailing the mode of action over gene expression produced by the application of these compounds (Arvanitidou et al., 2024). In this work, transcriptomic patterns in *Triticum aestivum (T. aestivum)* plants under conditions of drought stress and full irrigation were analyzed after foliar application of the microalgae-based biostimulant LRM™. Phenotypic effects were also determined and associated to changes in gene expression, to unveil the molecular basis of the product’s agronomic benefits.

## Materials and methods

### Plant growth and sample collection

*Triticum aestivum* seeds variety ‘Chinese Spring L42’ (WBCDB0017) were acquired from the Germplasm Resource Unit (GRU) at the John Innes Center, Norwich (UK). Seeds were stratified for 4 days at 4°C in the dark and subsequently germinated on plates in growth chambers under long summer day conditions (16h light: 8h dark) at 24°C. Seedlings were then planted in pots in the greenhouse, which had been pre-irrigated to their maximum retention volume of 180 ml. Plants were divided into four groups: fully irrigated control plants, fully irrigated treated plants, drought-stressed control plants and drought-stressed treated plants. Fully irrigated plants were watered twice a week with 180 ml and drought-stressed plants with 30 ml. The greenhouse conditions were long summer days, with 22°C during the day and 20°C during the night. The microalgae-based biostimulant LRM™ was provided by AlgaEnergy (Madrid, Spain). Following manufacturer’s instructions, LRM™ was diluted at 0.5% in water and was applied by foliar spraying over treated plants on the next day after planting and every 14 days thereafter. Control plants were sprayed with water.

Samples were collected two hours after the second application of the biostimulant 14 days after planting during the tillering stage. Three replicates were collected for each group of plants resulting in twelve samples. Each sample consisted in the last fully expanded leave of five different plants from the same group. Samples were immediately frozen in liquid N_2_ and stored at −80 °C until processing.

### RNA extraction, sequencing and analysis

For RNA extraction, plant samples were ground to a fine powder in liquid N_2_ and TRIsure™ (BIOLINE, Memphis, TN, USA) was used following manufacturer’s instructions. RNA purification was carried out using ISOLATE II RNA Plant Kit (BIOLINE) and RNA integrity was assessed with a Bioanalyzer Agilent 2100 system. Illumina’s instructions were followed for mRNA libraries preparation and sequencing was performed on a NextSeq500 Illumina sequencer (Illumina, San Diego, CA, USA) producing approximately 40 million reads of 100 nucleotides in length per sample. Quality control was performed with FASTQC. Short read mapping to the reference genome (Triticum 4.0 assembly GenBank accession: GCA_002220415.3) (Alonge et al., 2020) was carried out with HISAT2 (Kim et al., 2019). Transcripts assembly and gene expression estimation based on discrete mapped read counts was performed using StringTie (Kovaka et al., 2019). Differential gene expression was determined using the Bioconductor R package DESeq2 (Love et al., 2014) using a fold-change of 2 and an adjusted p-value of 0.05. For bar plot representations, gene expression was estimated as FPKM (Fragments Per Kilobase of exons and Millions of mapped reads). The R package FactoMineR was used for Principal Components Analysis and visualization (Lê et al., 2008).

### Functional Enrichment Analysis

An R package for Gene Ontology (GO) functional annotation, called ‘org.Taestivum.eg.db’ (https://greennetwork.us.es/) was developed by our group. The R package AnnotationForge and the functional annotation based on orthologous relationships with *Arabidopsis thaliana* developed in (Alonge et al., 2020) were used. The bioconductor R package clusterProfiler (Wu et al., 2021) was used to identify significantly enriched biological processes GO terms in the different gene sets of interest, using a Benjamini-Hochberg adjusted p-value of 0.05.

### Phenotypic analysis

Plants not used for sample collection were allowed to proceed through their entire life cycle following the previously described watering and treatment regimes. Spikes were collected, measured and classified into four groups: fully irrigated control, fully irrigated treated, drought-stressed control and drought-stressed treated. Data normality for spike length was assessed using the Shapiro–Wilk test, which rejected the null hypothesis of normality. Consequently, the Mann–Whitney-Wilcoxon test was applied to analyze significant differences between the rank of the values in each group.

## Results and Discussion

### Phosphate starvation response, inositol phosphate signaling and tocopherol biosynthesis are activated under drought stress in bread wheat

Our gene expression analysis found significant differences between the transcriptomes of fully irrigated (H_2_O 180 ml) and drought-stressed (H_2_O 30 ml) plants (**Fig. 1A**). A clear activation in gene expression was observed with 348 genes being significantly activated under drought conditions, when compared to full irrigation, whereas only 28 genes were significantly repressed (**Supplementary Table S1**). Functional enrichment analysis over the activated genes identified biological processes related to nutrient starvation, including phosphate, sulfur and nitrate deprivation, inositol metabolic process and tocopherol (vitamin E) biosynthesis (**Supplementary Table S2**). Specifically, a recurrent activation of processes related to cellular response to phosphate starvation was detected with processes such as phosphate ion homeostasis and glycerophospholipid catabolic process being significantly activated. This is in agreement with recent studies in the characterization of early drought response in soybean (*Glycine max*). A different mechanism from the one traditionally known involving abscisic acid (ABA) signaling was revealed encompassing the activation of genes associated with phosphate starvation response (PSR). A reduction in inorganic phosphate (Pi) availability was found to be concomitant to drought stress affecting cop growth (Nagatoshi et al., 2023). The PSR pathway is initiated by signaling through SPX receptors and PHR transcription factors, which modulate the expression of phosphate transporters and other genes involved in increasing intracellular Pi concentration, such as ribonucleases (RNases), phosphatases and lipid remodeling enzymes (Gho et al., 2020; Nagatoshi et al., 2023). Our analysis found a significant increase in expression in the genes coding for these PSR biomarker proteins in drought-stressed wheat plants. Activation was observed throughout the entire pathway, from genes codifying receptors and transcription factors, *SPX* and *PHR2,* phosphate transporters, *PHT*, to enzymes involved in lipid degradation, *SQD1*, *SQD2* and *MGD2*, ribonucleases *RNS2* and phosphatases *PAP10* (**Fig. 1B**).

**Figure 1.**
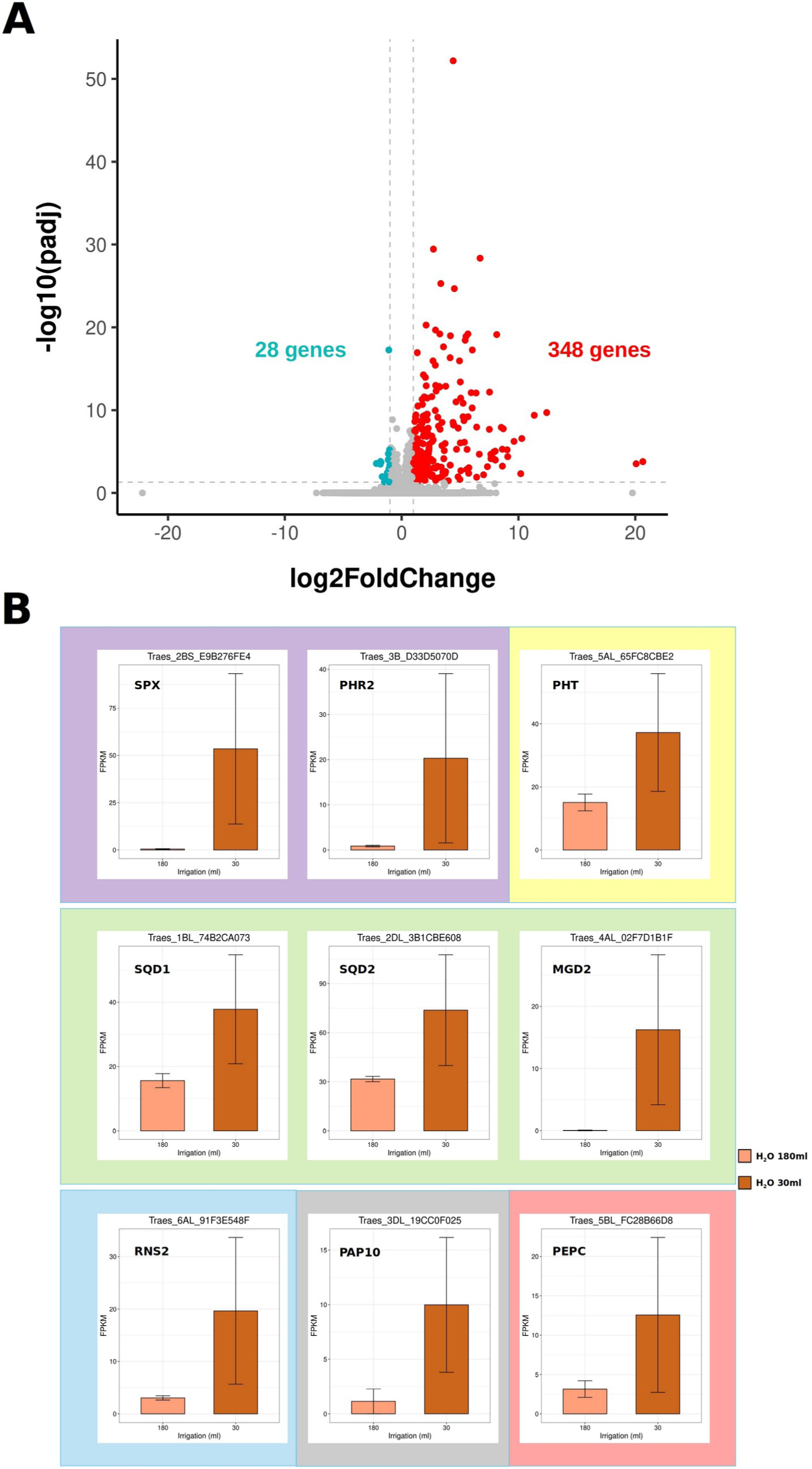
Wheat transcriptomic response to drought. **A**. Volcano plot showing activated (red), repressed (blue) and not significantly changed (gray) genes upon drought conditions. Fold change and significance thresholds are shown as dashed lines. **B**. Gene expression of PSR biomarkers in fully irrigated (light brown, H_2_0 180 ml) and drought-stressed (dark brown, H_2_0 30 ml) untreated wheat plants. Genes are classified into signaling (purple), phosphate transporters (yellow), lipid degradation (green), RNAses (blue), phosphatases (gray) and PEPC (red).

Additionally, genes encoding phosphoenolpyruvate carboxylase (PEPC) also showed strong activation in drought-stressed wheat plants. This enzyme has been previously associated with PSR, with increased gene expression and posttranslational modifications in situations reported under limiting phosphate conditions in the model plant *Arabidopsis thaliana* (Gregory et al., 2009) and in the leguminous plant *Sesbania rostrata* (Aono et al., 2001), among others. In the absence of Pi, PEPC not only generates Pi from phosphoenolpyruvate (PEP), but also mediates the synthesis of pyruvate without the use of adenine nucleotides, which are not readily available under these conditions (Nagano et al., 1994).

Our analysis revealed significant overexpression of four genes coding for IP6K/VIH in drought-stressed wheat plants, indicating an activated inositol metabolic process as identified in our functional enrichment analysis (**Supplementary Table S2**). Previous studies have reported raised levels of the inositol pyrophosphate InsP7 under low Pi concentrations (Williams et al., 2015), consistent with the overexpression of these genes identified in our study. These signaling molecules have been shown to be involved in responding to water stress by favoring plant survival through modulation of cell wall composition (Shukla et al., 2021). This suggests that the activation of inositol biosynthesis could provide wheat plants with signaling systems to elicit responses that enhance their survival when facing drought.

A significant activation in tocopherol (vitamin E) biosynthetic process was found among the significantly activated genes in drought-stressed plants (**Supplementary Table S2**). This compound is used in plants to quench reactive oxygen species (ROS) generated under various stresses such as drought, temperature and high light stresses (Mesa & Munné-Bosch, 2023). The enhancement of tocopherol biosynthesis has been demonstrated to infer drought tolerance in tobacco plants (Liu et al., 2008). Consequently, activated tocopherol biosynthesis as a response to drought could be part of the resistance mechanisms to overcome this stress in bread wheat plants.

Although only 28 genes were found significantly repressed in our analysis (**Supplementary Table S1**) some biological processes were significantly repressed, as the transcription by RNA polymerase II, which explained the high ratio of activated to repressed genes. Besides, a small number of genes related to chlorophyll biosynthesis also appeared repressed, hinting at a readiness to slow down the photosynthetic process in the presence of drought (**Supplementary Table S2**).

These results indicate that drought produces phosphate deficiency in wheat plants, which triggers PSR to resist the lack of this nutrient by mobilizing available Pi in the cells.

### The microalgae-based biostimulant enhances the response to drought by further activating PSR and carbon fixation while inhibiting light harvesting in photosynthesis

The foliar application of the microalgae-based biostimulant LRM^TM^ on bread wheat plants under drought conditions produced a drastic effect on their transcriptome. Drought-stressed treated plants (LRM 30 ml group) were compared to drought-stressed control untreated plants (H_2_O 30 ml) two hours after treatment. Our analysis identified more than 1500 differentially expressed genes (DEGs) (**Supplementary Table S3**). In contrast to the previous comparison between drought-stressed and fully irrigated untreated plants, in this case, the number of significantly activated and repressed genes was similar, 867 activated genes and 719 repressed genes (**Fig. 2A**). This suggests that LRM^TM^ foliar treatment under drought stress has a broad impact on gene expression, with its regulatory effects involving both gene upregulation and downregulation.

**Figure 2.**
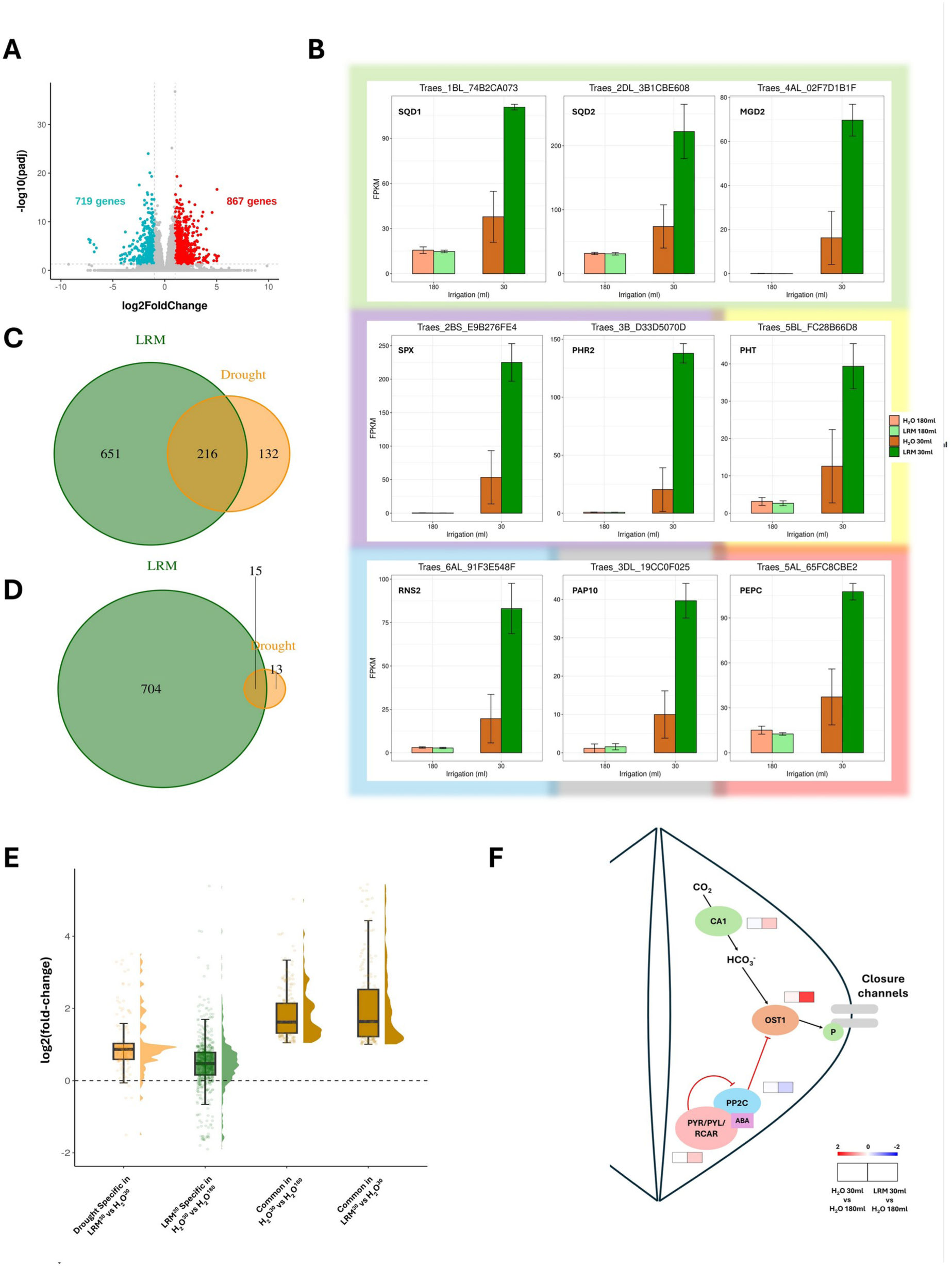
Transcriptomic response to LRM^TM^ foliar application in drought-stressed wheat plants. **A.** Volcano plot showing activated (red), repressed (blue) and not significantly changed (gray) genes two hours after LRM^TM^ foliar application in drought-stressed plants. **B.** Gene expression of PSR biomarkers in fully irrigated untreated (light brown, H_2_O 180 ml) and treated (light green, LRM 180 ml) plants, as well as, in drought-stressed untreated (dark brown, H_2_O 30 ml) and treated (dark green, LRM 30 ml) plants. Genes are classified into signaling (purple), phosphate transporters (yellow), lipid degradation (green), RNAses (blue), phosphatases (gray) and PEPC (red). **C.** Venn diagram showing the number of common and specific activated genes between the response to drought in untreated plants (brown) and the response to LRM^TM^ foliar application in drought-stressed plants (green). **D.** Venn diagram showing the number of common and specific repressed genes between the response to drought in untreated plants (brown) and the response to LRM^TM^ foliar application in drought-stressed plants (green). **E.** Raincloud plot showing the distribution log2-fold-changes or activation levels for drought specific activated genes in drought-stressed treated plants (LRM 30 ml vs H_2_O 30 ml contrast), LRM specific activated genes in drought-stressed untreated plants (H_2_O 30 ml vs H_2_O 180ml contrast), and common genes in both contrasts. **F**. ABA (bottom) and CO_2_ (top) pathways regulating stomata closure. Activation levels or log2-fold-changes are represented for each gene using red for activation and blue for repression. Left represents the response to drought in untreated plants and right corresponds to the combined effect of drought and LRM^TM^ treatment.

Functional enrichment analysis over the activated genes by LRM^TM^ under drought conditions revealed similar results to the ones produced only by drought in untreated plants including PSR as the most significant biological process affected (**Supplementary Table S4**). The expression levels of the genes coding for PSR biomarker proteins revealed a significant increase, after LRM^TM^ application, in drought-stressed wheat plants when compared to both, drought-stressed and fully irrigated untreated plants (**Fig. 2B**). Thus, the microalgae-based biostimulant enhanced the overexpression of specific drought-activated genes. In order to test whether this was a general trend not only restricted to PSR, the genes solely activated by drought in untreated plants were compared with the activated genes by LRM^TM^ treatment under drought conditions. This analysis revealed that 216 of the 348 genes (62%) activated by drought in untreated plants were further upregulated following the foliar application of the microalgae-based biostimulant (**Fig. 2C**). This indicates that the biostimulant not only increases the expression of PSR genes but also amplifies the overexpression of almost two thirds of the genes activated by drought stress.

The 651 genes that were specifically activated after LRM^TM^ treatment under drought conditions, but not by drought alone in untreated plants, were significantly involved in biological processes such as PSR, glutathione synthesis and protein modifications such as nitrosylation. Notably, these activated genes were also involved in carbon fixation. While the first three processes can be understood as typical responses to the adverse conditions associated to drought, the activation of carbon fixation genes by LRM^TM^ treatment is particularly intriguing. Recent studies have shown that short-term drought stress can promote photosynthetic carbon fixation, even though overall photosynthetic rates are typically reduced under such conditions (Wang et al., 2022). Additionally, a subset of 132 genes were specifically activated by drought in untreated plants but did not show further upregulation with LRM^TM^ treatment. These genes were primarily associated with PSR, vitamin E biosynthesis and phospholipid catabolism (**Supplementary Table S4**).

Functional analysis over the repressed genes by LRM^TM^ under drought conditions revealed enrichment in photosynthetic and chloroplast developmental processes, including chloroplast rRNA processing, chlorophyll biosynthetic process, photosynthesis, light harvesting and phylloquinone biosynthetic process (**Supplementary Table S4**). Phylloquinones are involved in electron transport through photosystem I and the creation of disulfide bridges between photosystem II subunits (Reumann, 2013). These molecules are predominantly localized in chloroplasts, and the repression of their biosynthesis, along with the downregulation of genes involved in plastid transcription, translation, localization, fission and membrane organization, suggests an inhibition of the photosynthetic process. This is further supported by the repression of processes such as the synthesis of chlorophylls and carotenoids. Downregulation of genes involved in light harvesting in photosynthesis under drought conditions is a known protective response to reduce the accumulation of ROS in chloroplasts (Pinheiro & Chaves, 2011). Therefore, the foliar application of the microalgae-based biostimulant enhanced this protective inhibition of light harvesting that was observed only in a priming stage in the comparison between drought-stressed and fully irrigated untreated plants where only chlorophyll synthesis was significantly repressed (**Supplementary Table S2**). This repression of light harvesting did not show any detrimental effect over the photosynthetic capacity or growth in treated plants.

Further analysis revealed that a significant proportion of drought-repressed genes in untreated plants were more strongly repressed following LRM^TM^ treatment, with 15 out of 28 genes (∼54%) exhibiting increased repression (**Fig. 2D**). These 15 genes are associated with the negative regulation of transcription by RNA polymerase II and hexose metabolism, in agreement with the observed extensive activation of stress-responsive genes and photosynthesis inhibition. However, a pronounced specific effect was observed with 704 genes (∼96%) specifically repressed by the biostimulant under drought conditions. Nevertheless, the same enriched biological processes were identified in this subset indicating again an enhanced response to drought induced by the foliar application of LRM^TM^.

To further explore this enhancement, the distributions of fold-changes or activation levels of genes commonly and specifically activated by LRM^TM^ under drought conditions, as well as those activated by drought alone, were analysed across all comparisons (**Fig. 2E**). Most of the genes activated solely by drought in untreated plants also presented positive fold-changes in the comparison between drought-stressed treated and untreated plants (LRM 30 ml versus H_2_O 30 ml), left most orange boxplot in **(Fig. 2E)**. However, these increases in expression were not substantial or significant enough according to the selected thresholds (see Materials and methods) to be considered as activated. This indicates that nearly all genes activated by drought were further upregulated upon LRM^TM^ treatment with variable levels of increase. In contrast, the genes exclusively activated by the foliar application of LRM^TM^ in drought-stressed plants exhibited both positive and negative fold-changes with median close to zero in the comparison between drought-stressed and fully irrigated untreated plants (H_2_O 30 ml versus H_2_O 180 ml), green boxplot in **(Fig. 2E)**. This shows a specific effect of the treatment with the biostimulant that cannot be elicited by drought stress alone. As expected, similar high fold-change values were present in the commonly activated genes in the corresponding contrasts, as seen in the brown boxplots in (**Fig. 2E**).

These results demonstrated a boosting effect of LRM^TM^ on almost all drought responsive genes, and the existence of specific genes activated by the biostimulant. Only a few of the LRM^TM^-specifically activated genes showed initial upregulation by drought stress.

### The microalgae-based biostimulant specifically activates ABA-based stress response leading to the activation of genes involved in stomata closure

As discussed above, the microalgae-based biostimulant LRM^TM^ has specific effects over the transcriptome of drought-stressed plants that are not upregulated during the early onset of drought stress in untreated plants. A more detailed analysis of these sets of LRM^TM^-specific genes revealed typical responses induced by ABA and CO_2_ signaling, particularly those involved in the regulation of stomata closure. Although, a clear photosynthetic inhibition was enhanced after LRM^TM^ foliar application by repressing specific genes, CO_2_ diffusion through stomata has been described as a crucial factor influencing photosynthesis activity (Pinheiro & Chaves, 2011). Furthermore, stomata closure regulation has been shown to play a critical role in improving drought tolerance in wheat plants as it reduces water loss through transpiration (Onyemaobi et al., 2021). Our transcriptomic analysis reflected an impact on the central regulator of stomatal closure OPEN STOMATA 1 (OST1). While, in drought-stressed and fully irrigated plants, *OST1* gene expression was practically unaffected, the biostimulant foliar application produced a strong upregulation in the expression of this gene. ABA signaling pathway is a key regulatory mechanism regulating *OST1* gene expression. ABA binds to PYRABACTIN RESISTANCE / PYRABACTIN RESISTANCE-LIKE / REGULATORY COMPONENT OF ABA RECEPTOR (PYR/PYL/RCAR) receptors, preventing PROTEIN PHOSPHATASE 2C (PP2C) phosphatases from inhibiting OST1. Another important mechanism regulating *OST1* expression is the CO₂-bicarbonate signaling pathway in which CO_2_ is transformed in the guard cells to HCO_3_^-^ by CARBONIC ANHYDRASE 1/4 (CA1/4). In turn, high HCO_3_^-^ concentrations favor OST1 activation. These conditions are favored by drought stress, which promotes stomatal closure through OST1 phosphorylation of anion channels, with an associated efflux of malate (Daloso et al., 2017).

Changes in the expression of genes involved in these two signaling pathways because of water deficiency alone (H_2_O 30 ml vs H_2_O 180 ml) and as a consequence of LRM^TM^ foliar application over drought-stressed plants (LRM 30 ml vs H_2_O 180 ml) are depicted in **(Fig 2F)**. In general, no significant gene expression change was observed induced solely by drought. However, LRM^TM^ treatment triggered a response comprising both pathways. On the one hand, a clear *CA1* overexpression was observed which would produce an increase in HCO ^-^ levels activating *OST1* expression. On the other hand, in the ABA signaling pathway, an activation in *PYL7* gene expression coupled with a repression in *PP2C* gene expression was observed, which would further contribute to the activation of *OST1*. These results indicate that LRM^TM^ application is specifically inducing stomata closure, thereby enhancing drought tolerance in treated plants. This response is absent in control untreated wheat plants during the early stages of drought stress.

Recent studies have shown that mild drought stress initially triggers PSR-based responses, whereas more severe or progressive stress activates ABA signaling to cope with the associated deleterious effects (Nagatoshi et al., 2023). Our study, comparing drought-stressed and fully irrigated untreated plants, is in agreement with these results, particularly in the overexpression of PSR genes at the onset of drought stress. Additionally, our analysis revealed upregulation of the expression of ABA-responsive genes as a specific LRM^TM^ effect under drought conditions. This suggests that LRM^TM^ treatment accelerates and amplifies drought response, shifting it from PSR to ABA signaling. Therefore, the microalgae-based biostimulant enhances drought tolerance in treated plants by reinforcing their response to water deficiency.

### LRM^TM^ foliar application over fully irrigated plants reduces stress responses by modulating shade response, circadian rhythms and endomembrane trafficking

The effect of the microalgae-based biostimulant LRM^TM^ over the transcriptome of fully irrigated (180 ml) wheat plants was also analysed. The impact was considerably smaller compared to the effect observed in drought-stressed plants with only 35 activated and 25 repressed genes (**Fig. 3A, Supplementary Table S5)**. Recall that our analysis included twelve different samples, three replicates for each one of the four conditions under study: fully irrigated untreated (H_2_O 180 ml), drought-stressed untreated (H_2_O 30 ml), fully irrigated LRM^TM^ treated (LRM 180 ml) and drought-stressed LRM^TM^ treated (LRM 30 ml). Principal component analysis (PCA) of these samples revealed separate and distinct clusters for LRM^TM^ treated and untreated plants under drought, whereas the samples corresponding to fully irrigated plants, both treated and untreated grouped together (**Fig. 3B**). These results indicate that the microalgae-based biostimulant exerts a strong transcriptomic effect during the tillering stage in drought-stressed plants, whereas only a slight effect was detected in fully irrigated plants.

**Figure 3.**
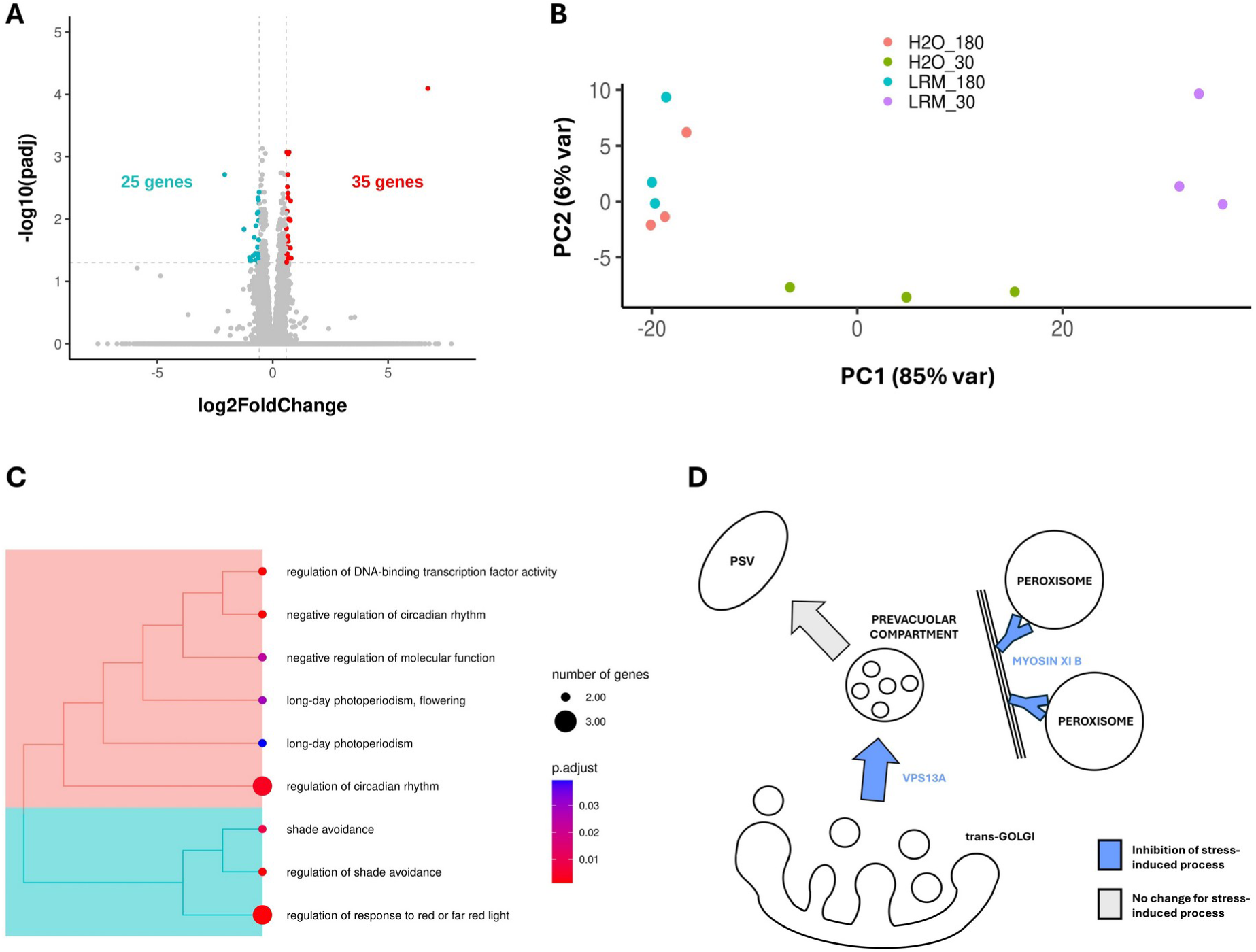
Transcriptomic response to LRM^TM^ foliar application in fully irrigated wheat plants. **A.** Volcano plot showing activated (red), repressed (blue) and not significantly changed (gray) genes upon LRM^TM^ application in fully irrigated plants. **B.** Principal Component Analysis of samples corresponding to fully irrigated untreated plants (H_2_O 180 ml, brown), drought-stressed untreated plants (H_2_O 30 ml, green), fully irrigated treated plants (LRM 180 ml, blue) and drought-stressed treated plants (LRM 30 ml, purple). **C.** Significantly enriched GO terms in activated genes upon LRM^TM^ application under full irrigation. Blue: processes related to circadian rhythm, red: processes related to shade avoidance. **D.** Pathways regulating trafficking in endomembranes systems as a response to stress. Blue represents inhibition and gray no significant change as detected in fully irrigated wheat plants after LRM^TM^ treatment.

Functional analysis over the activated genes by LRM^TM^ under fully irrigation revealed significant enrichment in processes related to circadian rhythm regulation, shade avoidance and response to red and far-red light (**Fig. 3C, Supplementary Table S6**). These processes are traditionally of agronomic interest due to their relationship with crop efficiency (Bendix et al., 2015; Tang & Liesche, 2017) suggesting beneficial effects of the biostimulant also in non-stressed plants.

Specifically, LRM^TM^-treated fully irrigated plants exhibited increased gene expression levels for CIRCADIAN CLOCK ASSOCIATED 1 (CCA1), a transcription factor central to circadian rhythms regulation. Overexpression of *CCA1* has been shown to attenuate ROS stress induced by high midday irradiance (Lai et al., 2012; Nagano et al., 1994), regulate stomatal aperture, increasing leaf size and biomass values under well irrigated conditions (Hassidim et al., 2017), and affect wheat agronomic properties (Gong et al., 2022).

In our functional analysis over repressed genes (**Supplementary Table S6**), deadenylation-dependent decay of mRNA was identified as a significantly enriched process, with repressed genes such as CARBON CATABOLITE REPRESSION 4-NEGATIVE ON TATA-LESS (CCR4-NOT). This mechanism has been linked to stress responses (Walley et al., 2010), and its repression could point to a biostimulant-mediated relaxation of stress responses in their absence. CCR4-NOT has also been described as a regulator of phytochrome A-mediated signaling (Schwenk et al., 2021; Shen et al., 2009), one of the key players in the response to red and far-red light detected among the activated genes. Processes involving protein localization to vacuoles, Golgi apparatus and peroxisomes were also significantly repressed. Abiotic stresses have a substantial impact on the endomembrane system, rearranging its trafficking to favor the accumulation of proteins and other compounds in specific organelles (Neves et al., 2021; Sampaio et al., 2022). Under stress conditions, two trafficking pathways, among others, have been described to be upregulated; the trans-Golgi network to the prevacuolar compartment, followed by the protein storage vacuole pathway mediated by proteins such as VPS13 (VACUOLAR PROTEIN SORTING-ASSOCIATED PROTEIN 13), and the accumulation of peroxisomes, through myosin XI-B (Hickey et al., 2022; Hinojosa et al., 2019; Neves et al., 2021; Tian et al., 2021). Our analysis showed repression of both processes with reduced expression levels of genes encoding VPS13 and myosin XI-B in fully irrigated treated plants suggesting a biostimulant-mediated repression of abiotic stress responses (**Fig. 3D**).

No significant overlap was detected between genes activated or repressed by LRM^TM^ under drought conditions and those modified under fully irrigation. This indicates that the biostimulant employs distinct pathways to modulate gene expression in the different conditions.

### Phenotypic effect of LRM^TM^ treatment under Drought Stress and Full Irrigation

To complement this study, the phenotypic effect of the microalgae-based biostimulant was evaluated across the entire life cycle of drought-stressed and fully irrigated wheat plants. This analysis included the tillering, stem elongation and ripening stages focusing on the number of tillers, plant height and spike length.

Our transcriptomic analysis was conducted two hours after the second foliar application of the LRM^TM^ biostimulant which coincided with the tillering stage. At this point, the effects of drought stress began to manifest, with fully irrigated plants exhibiting more tillers than drought-stressed plants, as shown in the lower and upper left panels in (**Fig. 4**). Although differences between treated (LRM^TM^, left) and untreated (H_2_O, right) plants were small, a slight increase in tiller number was observed in treated plants.

**Figure 4.**
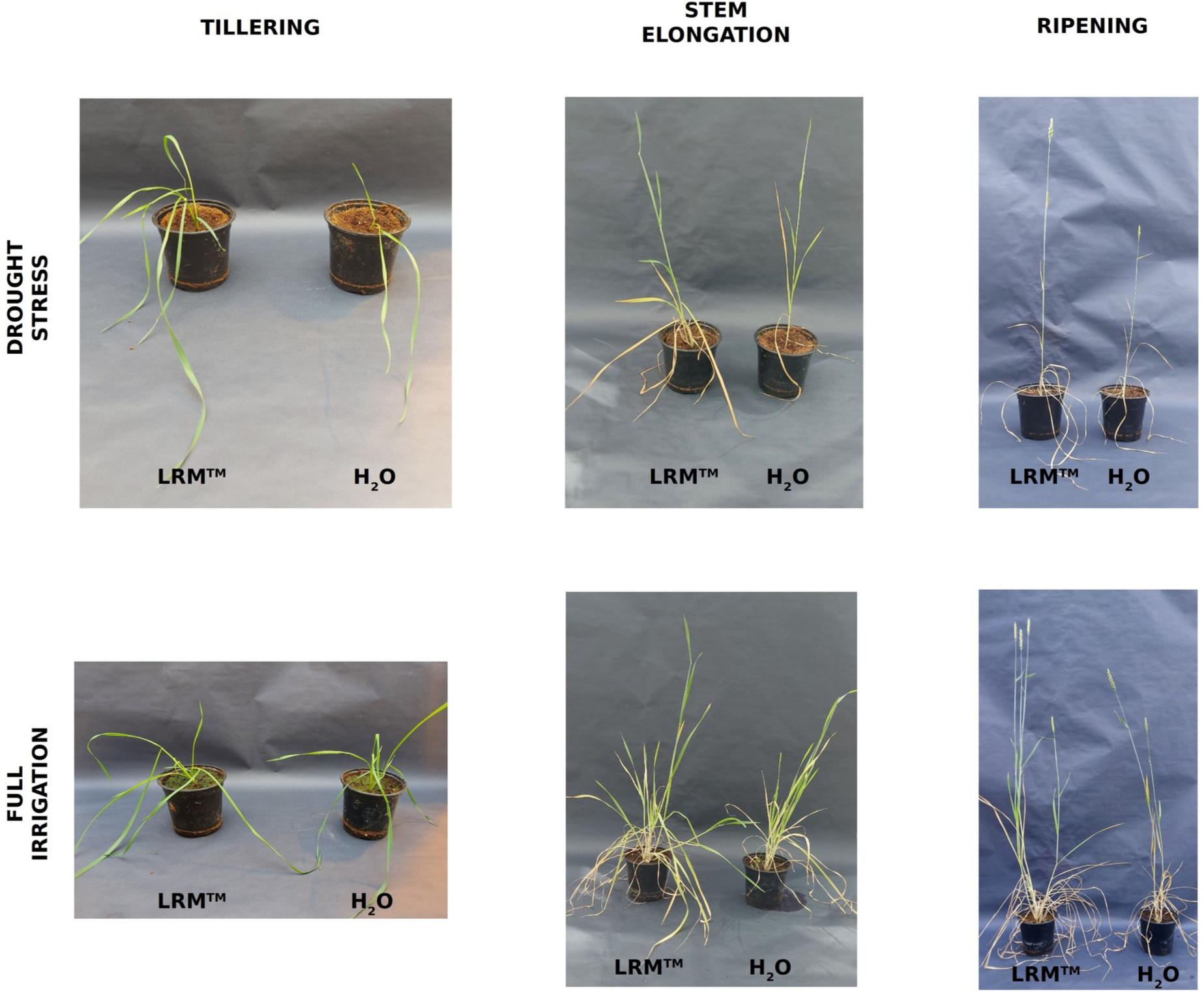
Phenology of LRM^TM^ treated and untreated plants under drought stress and full irrigation at tillering, stem elongation and ripening developmental stages.

Plants continued to be treated with LRM^TM^ biostimulant every 14 days while untreated plants were sprayed with water. During the stem elongation stage, drought-stressed plants, both treated and untreated, primarily developed a single stem, whereas fully irrigated plants exhibited multiple stems. LRM^TM^ treated plants under both irrigation regimes displayed greater height compared to untreated plants, as depicted in the central panels in (**Fig. 4**). This observation is in agreement with transcriptomic data, which indicated significant activation of shade avoidance and enhanced drought response in treated plants. Our results are consistent with previous studies on the effect of drought in wheat, reporting a decrease in the number of tillers and height in stressed plants. These reductions were less pronounced in wheat genotypes that are more tolerant to drought (Poudel et al., 2020).

Ripening was the last stage analysed in this study. Most drought-stressed plants developed a single spike, whereas fully irrigated plants produced multiple spikes, due to the presence of several stems. Treated plants consistently exhibited more stems and, consequently, more spikes, as shown on the right panels in (**Fig. 4**). The greater height observed in treated plants persisted during the entire life cycle corroborating transcriptomic analysis. These results suggest that the microalgae-based biostimulant continued enhancing drought response in stressed plants and inferring beneficial features in fully irrigated plants over the entire life cycle of the plants.

Spike lengths were measured as an indicator of crop yield for fully irrigated treated (**Fig. 5A**) and untreated (**Fig. 5B**) plants, as well as in drought-stressed treated (**Fig. 5C**) and untreated (**Fig. 5D**) plants. As expected, spikes were significantly shorter in drought-stressed plants compared to fully irrigated plants (Frantová et al., 2022). However, plants treated with the microalgae-based biostimulant under both irrigation regimes developed significantly longer spikes (p-value < 10^-3^), resulting in higher yield compared to untreated plants. The beneficial effect was more pronounced in drought-stressed plants, where treated plants presented spikes that were 41% longer than those of untreated plants, whereas the spikes of fully irrigated treated plants were 16% longer than those of untreated plants (**Fig. 5E**).

**Figure 5.**
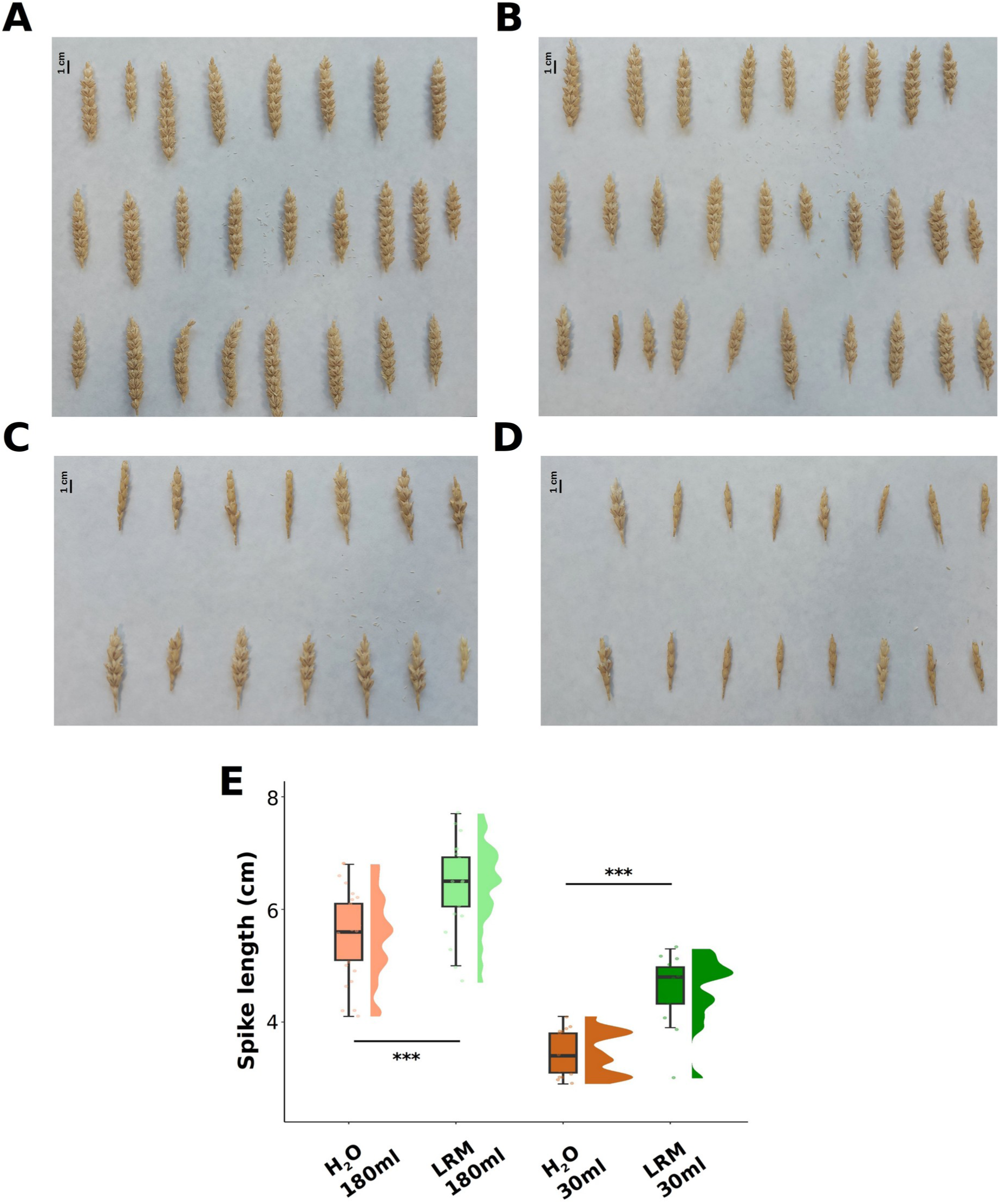
Effect of LRM^TM^ treatment over spike length. **A.** Spikes from fully irrigated treated plants. **B.** Spikes from fully irrigated untreated plants. **C.** Spikes from drought-stressed treated plants. **D.** Spikes from drought-stressed untreated plants. **E.** Raincloud plot showing significant differences between spike length distributions in fully irrigated untreated plants (light brown, H_2_O 180 ml), fully irrigated LRM^TM^ treated plants (light green, LRM 180 ml), drought-stressed untreated plants (dark brown, H_2_O 30 ml) and drought-stressed treated plants (dark green, LRM 30 ml).

## Conclusions

This study aimed to address the gap in available transcriptomic analysis concerning the effects of microalgae-based biostimulants on crops by demonstrating the positive impact of the biostimulant LRM™ (AlgaEnergy, Madrid, Spain) on bread wheat under both full irrigation and drought conditions. Foliar treatment of stressed plants led to an acceleration and amplification of drought responses, upregulating not only drought stress-related genes, including those involved in phosphate starvation response, but also biostimulant-specific genes, notably those involved in stomata closure. In fully irrigated plants the biostimulant also had a positive effect by downregulating stress response processes. These results, along with phenotypic observations, highlight the beneficial effects of the microalgae-based biostimulant LRM™ and pave the way for further research in sustainable agriculture.

## Author contributions

Conceptualization, M.G.G. and F.J.R.C.; methodology, C.A., M.E.G.G., M.R.G, M.G.G, and F.J.R.C.; software, C.A., M.R.G, and F.J.R.C; validation, C.A., M.E.G.G., M.R.G, M.G.G, and F.J.R.C.; formal analysis, C.A., M.R.G, and F.J.R.C; investigation, C.A., M.E.G.G., M.R.G, M.G.G and F.J.R.C.; resources, M.G.G. and F.J.R.C.; data curation, C.A., M.R.G, and F.J.R.C.; writing—original draft preparation, F.J.R.C., M.R.-G. and M.G.-G.; writing—review and editing, F.J.R.C., M.R.G. and M.G.G.; visualization, F.J.R.C., M.R.G. and M.G.G.; supervision, F.J.R.C. and M.G.G.; project administration, M.G.G. and F.J.R.C.; funding acquisition, M.G.G. and F.J.R.C. All authors have read and agreed to the published version of the manuscript.

## Acknowledgements

We thank Eloisa Andújar-Pulido, Mónica Pérez, Victoria Jiménez-Santos and Lola Pérez, from the Genomics Unit (CABIMER), for their assistance in RNA-sequencing data generation. We would also like to thank Miguel G. Guerrero for critically reading this manuscript. This work was supported by the research projects MICROCLIMATT (O00000226E2000044796) from the Spanish Ministry of Agriculture, Fisheries and Food, ELECTRA (PID2021-1210) and PERSEPHONE (PID2024-158798OB-I00) from the Spanish Ministry of Science and Innovation. CA was supported by Junta de Andalucía predoctoral grant PREDOC_00999 and MRG was supported by an FPU predoctoral grant FPU22/00511 from the Spanish Ministry of Science and Innovation.

## Data availability statement

RNA-seq data generated in this study is freely available from the Gene Expression Omnibus (GEO) database under the accession number GSE275202. The data analysis code developed using the statistical programming language R is freely available from the following GitHub repository TORRID: https://github.com/fran-romero-campero/TORRID.

## Supporting information

**Supplementary Table 1.** Differentially expressed genes in drought-stressed plants compared to fully irrigated plants.

**Supplementary Table 2.** Significantly enriched biological processes in the differentially expressed genes in drought-stressed plants compared to fully irrigated plants.

**Supplementary Table 3.** Differentially expressed genes in drought-stressed LRM^TM^ treated plants compared to drought-stressed untreated plants.

**Supplementary Table 4.** Significantly enriched biological processes in the differentially expressed genes in drought-stressed LRM^TM^ treated plants compared to drought-stressed untreated plants.

**Supplementary Table 5.** Differentially expressed genes in fully irrigated LRM^TM^ treated plants compared to fully irrigated untreated plants.

**Supplementary Table 6.** Significantly enriched biological processes in the differentially expressed genes in fully irrigated LRM^TM^ treated plants compared to fully irrigated untreated plants.

